# Autoinducer-2 functions as both a quorum sensing and metabolic signal in *Escherichia coli*

**DOI:** 10.64898/2026.06.09.730415

**Authors:** Noémie Godrie, Xuanlin Chen, Nicolas Näpflin, Lukas Malfertheiner, Assa Yeroslaviz, Christopher Schubert, Rin Ho Kim, Christian von Mering, Leanid Laganenka

## Abstract

Bacteria integrate diverse environmental signals to coordinate behavior, yet the relationship between nutrient sensing and quorum sensing (QS) remains incompletely understood. Autoinducer-2 (AI-2) is unique among QS signals in that its production is tightly linked to central metabolism, raising fundamental questions about the boundary between metabolic and signaling functions. In *Escherichia coli*, AI-2 coordinates collective behaviors through the *lsr* operon, whose expression is controlled not only by the AI-2-responsive repressor LsrR but also by the cAMP receptor protein (CRP), placing it at the intersection of carbon sensing and population-level signaling. While inhibition of *lsr* operon expression by PTS sugars such as glucose was previously established, we demonstrate that non-PTS sugars similarly suppress *lsr* expression through CRP, further decoupling QS activation from cell density and coupling it to carbon source availability. Systematic analysis of *Enterobacteriaceae* genomes reveals that CRP binding sites in the *lsr* promoter region are broadly conserved, indicating that metabolic modulation of AI-2 signaling is an ancestral regulatory feature. Importantly, using a FRET-based biosensor, we show that AI-2 uptake modulates intracellular cAMP levels in a manner resembling non-PTS carbon source transport, suggesting that AI-2 may have originally functioned as a nutrient substrate, with its signaling role emerging subsequently or co-evolving alongside. In support of this hypothesis, we isolated soil- and phyllosphere-associated bacteria capable of utilizing AI-2 as a sole carbon source. Our findings reveal an underappreciated metabolic dimension of AI-2 QS and suggest an evolutionary trajectory in which AI-2 signaling emerged from ancestral carbon utilization pathways.

**Importance:** QS allows bacteria to coordinate collective behaviors by detecting secreted signaling molecules, yet the evolutionary origins of these systems remain poorly understood. AI-2, one of the most broadly conserved bacterial signals, is derived from central metabolism and processed by machinery in *E. coli* that strikingly resembles a sugar utilization system. Here, we show that nutrient availability overrides cell density as the primary determinant of AI-2 responsiveness, that this regulatory logic is conserved among *Enterobacteriaceae* genomes, and that environmental bacteria can grow on AI-2 as a sole carbon source. These findings reframe AI-2 as a signal embedded within, and potentially evolved from, nutrient sensing pathways, with direct implications for understanding how byproducts of cellular metabolism can acquire signaling functions.

## Introduction

In natural, complex environments, bacteria continuously integrate multiple intra- and extracellular signals to adjust their physiological state in response to changing conditions (1-4). Key among these signals are nutrient cues, which inform cells about resource availability and shape global regulatory responses (5). Additionally, population density is monitored by many bacterial species to coordinate collective behaviors through a process known as quorum sensing (QS) (6). QS enables cells to produce, release, and detect small signaling molecules, thereby synchronizing gene expression across the population. This coordination allows bacteria to engage in behaviors that are energetically costly for individual cells but beneficial when performed collectively. Such behaviors include biofilm formation, secretion of public goods such as exoenzymes, and the production of virulence factors (6-8).

Among the many QS systems described in bacteria, autoinducer-2 (AI-2) is unusual in two respects. First, it is among the most phylogenetically widespread, being produced by both Gram-positive and Gram-negative species that encode the biosynthetic enzyme LuxS (9). Second, AI-2 is not a dedicated signaling molecule, but rather a byproduct of the activated methyl cycle, arising during the detoxification of S-adenosylhomocysteine (10, 11). Although the role of AI-2 as a signaling molecule versus a metabolic byproduct was long debated, it is now well established that AI-2 functions as a QS signal in many bacteria. This signaling is mediated by several receptor systems, including LsrB (e.g., in *E. coli* and *Salmonella*), LuxP (in *Vibrio* species), and more recently described dCACHE domains (12-14). In *Escherichia coli*, AI-2 regulates autoaggregation, biofilm formation, mammalian gut colonization, and interspecies interactions (15-22). Most of these phenotypes are regulated by chemotaxis towards AI-2, which is mediated by the LsrB AI-2-binding protein together with the Tsr chemoreceptor (20).

In *E. coli*, LsrB is encoded within the *lsr* operon, which encodes an ABC transporter for high-affinity AI-2 import, a kinase (LsrK) that phosphorylates intracellular AI-2, and the transcriptional repressor LsrR, which is inactivated by phospho-AI-2 to create a positive feedback loop driving operon expression (9). Notably, the *lsr* operon resembles a canonical carbohydrate utilization operon and is regulated not only by LsrR but also by the cAMP receptor protein (CRP), making it subject to carbon catabolite repression (CCR) (23-26). CCR operates through the phosphoenolpyruvate (PEP):phosphotransferase system (PTS), in which the phosphorylation state of the carrier protein EIIA^Glc^ reflects glucose availability. In the presence of glucose, dephosphorylated EIIA^Glc^ inhibits adenylate cyclase (CyaA), reducing cAMP production and thereby limiting CRP activity (27). The cAMP-CRP complex serves as a key transcriptional activator of over 180 genes in *E. coli*, most of which are involved in the metabolism of carbohydrates not transported by the PTS (28). Regulation of the central AI-2 QS operon by cAMP-CRP is therefore intriguing, as it suggests that the ability of *E. coli* to respond to cell density may be influenced by the availability of preferred carbon sources.

In this study, we provide further evidence for the integration of metabolic and quorum sensing regulatory systems in *E. coli*. We show that uptake of several PTS and non-PTS sugars inhibits *lsr* operon expression, further decoupling QS activation from cell density, whereas pyruvate, TCA cycle intermediates, and amino acids have no such inhibitory effect. Like PTS sugars, non-PTS sugar-mediated regulation occurs via cAMP-CRP, which acts as a transcriptional activator of the operon, while LsrR modulates its activity in response to extracellular AI-2. Analysis of CRP-binding sites across 19,795 *Enterobacteriaceae* genomes suggests this regulatory architecture is widespread and evolutionarily conserved. Notably, AI-2 import itself is associated with changes in intracellular cAMP levels, similarly to non-PTS carbon sources.

These observations raise the possibility that AI-2 may have originally functioned as a carbon source, either predating or co-evolving with its signaling role. Supporting this hypothesis, we isolated environmental bacterial strains capable of utilizing AI-2 as a sole carbon source. Taken together, our findings reveal an underappreciated metabolic basis of AI-2 quorum sensing and suggest that, in some species, nutrient utilization and intercellular signaling may have diverged from a common ancestral pathway.

## Results

### Both PTS and non-PTS sugars inhibit *lsr* operon expression

The mammalian gut provides a chemically complex and spatially structured environment in which free monosaccharides are present at high micromolar to millimolar concentrations and play a key role in driving *Enterobacteriaceae gut* establishment (29-31). To analyze the effect of different mammalian gut-derived carbon sources and metabolites on *lsr* operon expression, we incubated exponentially growing *E. coli* MG1655 cells carrying a plasmid-based P*lsr–gfp* transcriptional fusion with 1 mM of the PTS sugar glucose; the non-PTS sugars D-galactose, D-ribose, L-arabinose, and lactose; as well as pyruvate, acetate, succinate, and L-serine for 1 h. The levels of *lsr* operon expression were then assessed by measuring GFP fluorescence using flow cytometry.

Consistent with previous reports, incubation with glucose resulted in decreased *lsr* operon expression (Fig. 1A) (23). Similarly, and in line with observations for glycerol, all tested non-PTS sugars also inhibited *lsr* expression. Notably, the optical density of the cultures either significantly increased (e.g., in the presence of glucose) or was not affected by the presence of the substrates (Fig. 1B). These observations further support the notion that AI-2 quorum sensing is tightly integrated with central metabolism, and, more importantly, that nutrient availability can decouple AI-2 QS activity in *E. coli* from its cell density. In contrast to sugar treatments, incubation with gluconeogenic substrates, such as pyruvate, TCA cycle intermediates and L-serine, did not significantly affect *lsr* operon expression.

**Fig. 1.**
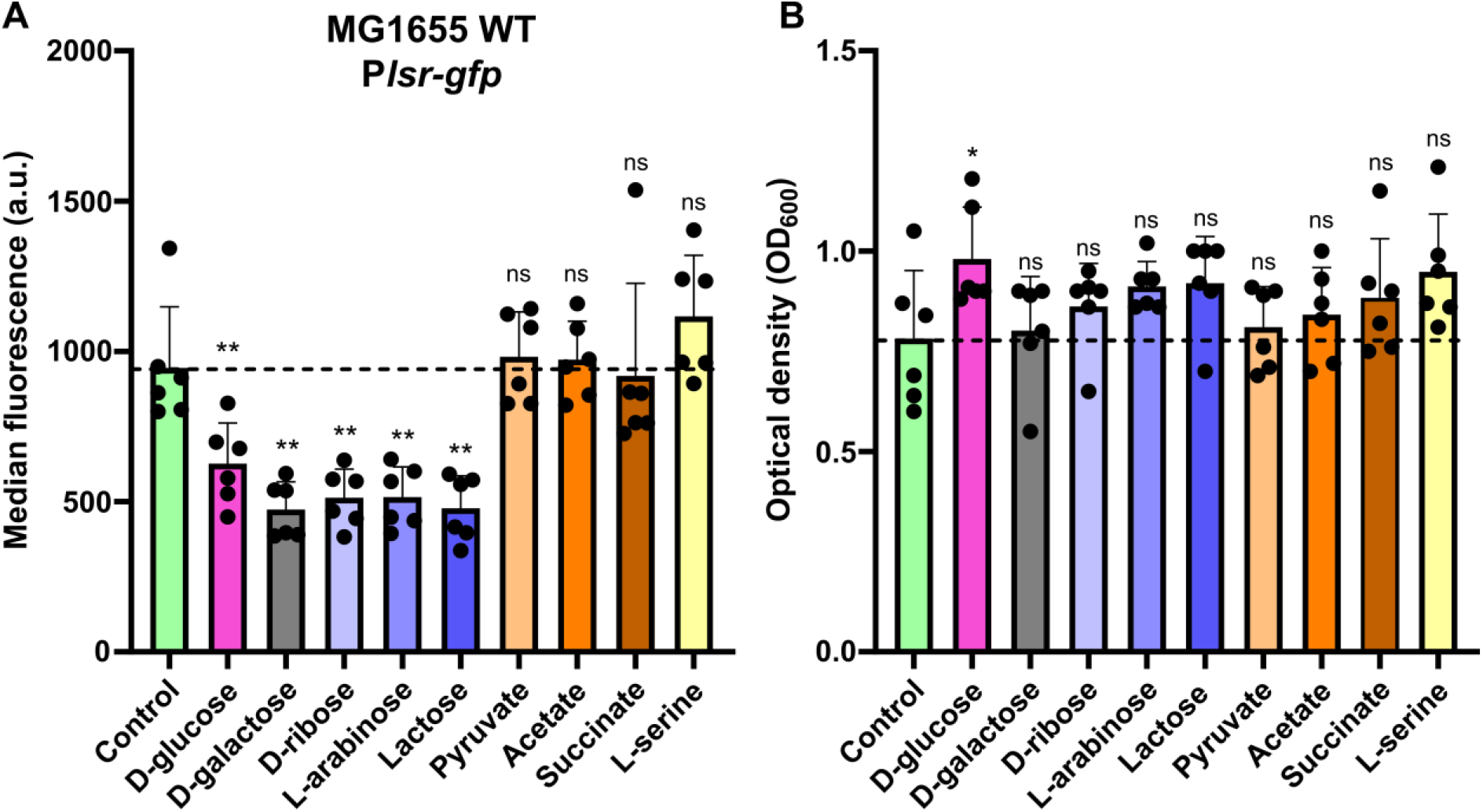
Both PTS and non-PTS sugars inhibit *lsr* operon expression in *E. coli* MG1655. **(A)** *lsr* promoter activity, measured by the fluorescence of a plasmid-based P*lsr-gfp* transcriptional reporter in the presence of 1 mM of the tested sugars (D-glucose, D-galactose, D-ribose, L-arabinose, lactose) and gluconeogenic substrates (pyruvate, acetate, succinate, and L-serine). **(B)** Optical density of *E. coli* MG1655 cultures shown in panel A. The dashed line represents the mean value of the control. *P* values were calculated using a two-tailed Mann-Whitney *U*-test (**P<0.01; ns, not significant). Bars indicate mean values (n=6, from two independent experiments), and error bars represent the standard deviation.

### cAMP-CRP is required for substrate-dependent modulation of *lsr* operon activity

Since glucose- and glycerol-dependent repression of the *lsr* operon has previously been shown to involve CRP, we hypothesized that the observed *lsr* operon inhibition might also occur via the same regulatory pathway. Although non-PTS sugars do not directly engage the PTS, their metabolism can still indirectly influence the phosphorylation state of PTS components and cellular energy status via PEP/pyruvate ratio sensing and concomitant changes in EIIA^Glc^ phosphorylation levels, thereby modulating cAMP levels and CRP activity (32).

Consistent with this hypothesis, deletion of *crp* resulted in a drastic reduction of basal *lsr* operon expression that could not be restored by addition of extracellular AI-2 (Fig. 2A, S1A). Moreover, inhibition of *lsr* operon activity by both PTS and non-PTS sugars was abolished in the Δ*crp* strain. Complementation of the Δ*crp* strain with a plasmid-borne wild-type *crp* allele restored sugar-dependent repression (Fig. 2B, S1B). Together, these results demonstrate that both PTS- and non-PTS sugar-dependent inhibition of *lsr* operon activity is mediated via the cAMP-CRP regulatory system, which acts as a transcriptional activator of the operon.

**Fig. 2.**
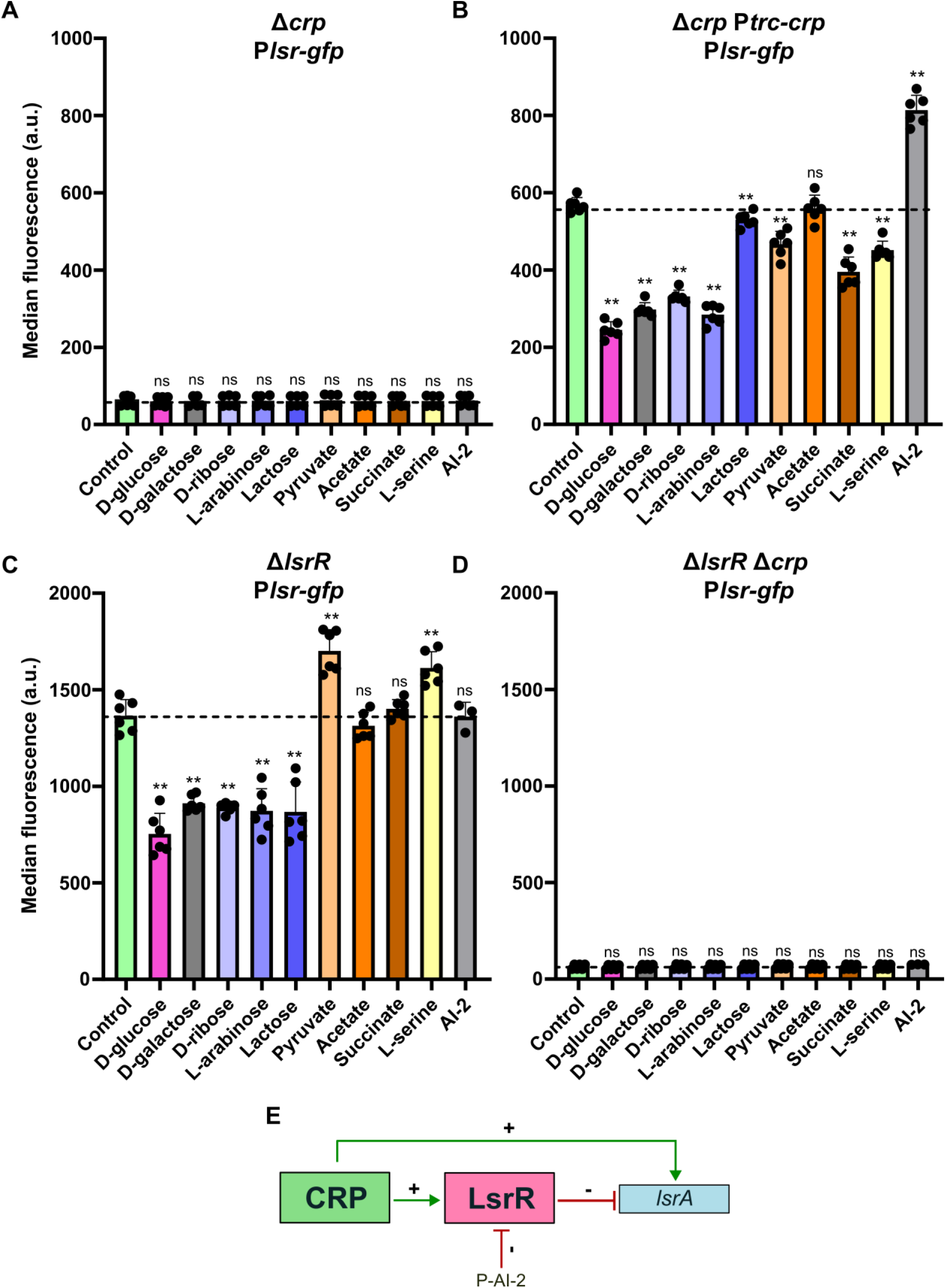
cAMP–CRP is required for substrate-dependent modulation of *lsr* operon activity. **(A)** *lsr* operon activity (P*lsr-gfp*) in the *E. coli* MG1655 Δ*crp* mutant in the presence of sugars, gluconeogenic substrates, and synthetic DPD/AI-2. **(B)** *lsr* operon activity (P*lsr-gfp)* in *E. coli* MG1655 Δ*crp* mutant strain complemented with a wild-type *crp* allele on the pTrc99a plasmid. Cells were grown in the presence of 7.5 µM IPTG to induce *crp* expression. **(C)** *lsr* operon activity in *E. coli* MG1655 Δ*lsrR* mutant and **(D)** in the Δ*lsrR* Δ*crp* double mutant in the presence of sugars, gluconeogenic substrates, and synthetic DPD/AI-2. The dashed line represents the mean value of the control. *P* values were calculated using a two-tailed Mann-Whitney *U*-test (**P<0.01; ns, not significant). Bars indicate mean values (n=6, from two independent experiments), and error bars represent the standard deviation. **(E)** Proposed model of a coherent CRP feed-forward regulatory loop. CRP directly activates transcription of both the *lsr* operon (*lsrA*) and *lsrR*, while LsrR acts as a repressor of the *lsr* operon. Under glucose-rich conditions, reduced intracellular cAMP levels limit CRP activity, leading to decreased expression of both *lsrA* and *lsrR*. In contrast, when alternative carbon sources are present, increased cAMP levels enhance CRP activity, promoting transcription of *lsrA* and *lsrR* and enabling LsrR-dependent regulation of the operon in response to phosphorylated AI-2 (P-AI-2).

To dissect the relative contributions of CRP and LsrR, we examined *lsr* expression in a Δ*lsrR* strain. At low cell density, when extracellular AI-2 concentrations are minimal, LsrR binds the *lsr* promoter and represses transcription. As extracellular AI-2 accumulates, it is imported by a yet unidentified low-affinity transporter, phosphorylated by LsrK, and phospho-AI-2 binds LsrR, relieving repression and allowing induction of the high-affinity import machinery (LsrACDB) and degradation enzymes (LsrFG) (9, 33, 34). This creates a positive feedback loop that amplifies *lsr* expression as AI-2 levels rise. As expected, basal *lsr* expression was elevated in the Δ*lsrR* strain, yet sugar-dependent inhibition was fully maintained. This demonstrates a hierarchical interplay between global and specific transcriptional control, in which cAMP-CRP provides the primary transcriptional drive while LsrR fine-tunes expression locally in response to extracellular AI-2 (Fig. 2C, S1C). Consistent with this, *lsr* operon activity in the Δ*lsrR* Δ*crp* double mutant was comparable to that in the Δ*crp* strain alone (Fig. 2D, S1D), confirming that CRP acts upstream of and independently from LsrR.

Interestingly, in the Δ*lsrR* background, incubation with pyruvate and L-serine led to an increase in *lsr* operon expression compared to the control. This effect was similarly dependent on CRP, as the stimulatory effect of pyruvate and L-serine was abolished in the Δ*lsrR* Δ*crp* strain (Fig. 2C, D). This suggests that gluconeogenic substrates may weakly promote cAMP-CRP activity under these conditions, though the mechanism and physiological relevance of this effect remain to be determined.

Taken together, these data indicate a coherent CRP feed-forward loop in which CRP directly activates both *lsrACDBFG* and *lsrR* expression, whereas LsrR regulates only the *lsr* operon, highlighting that functional cAMP-CRP signaling is essential for *lsr* expression (Fig. 2E) (23, 35). In the presence of glucose, low cAMP levels limit CRP activity and reduce transcriptional activation of the *lsr* operon and *lsrR*. Conversely, when alternative carbon sources are available, elevated cAMP levels promote cAMP-CRP-dependent activation of *lsr* and *lsrR*, enabling LsrR-mediated regulation of the *lsr* operon in response to AI-2.

### CRP binding sites in the *lsrA-lsrR* intergenic region are conserved among *Enterobacteriaceae*

To determine whether CRP regulation of the *lsr* operon is unique to *E. coli* or widespread across *Enterobacteriaceae*, we analyzed 19,795 genomes (Supplementary Table 3) for the presence of the *lsr* operon and scanned identified operons for putative CRP binding sites using a globally constructed search motif (Fig. S2, see Methods). The *lsr* operon was detected in 75% of all genomes analyzed (14,923 out of 19,795; Fig. 3), with no homologs identified in the enterobacterial genera *Cronobacter, Kosakonia*, or *Plesiomonas*.

**Fig.3.**
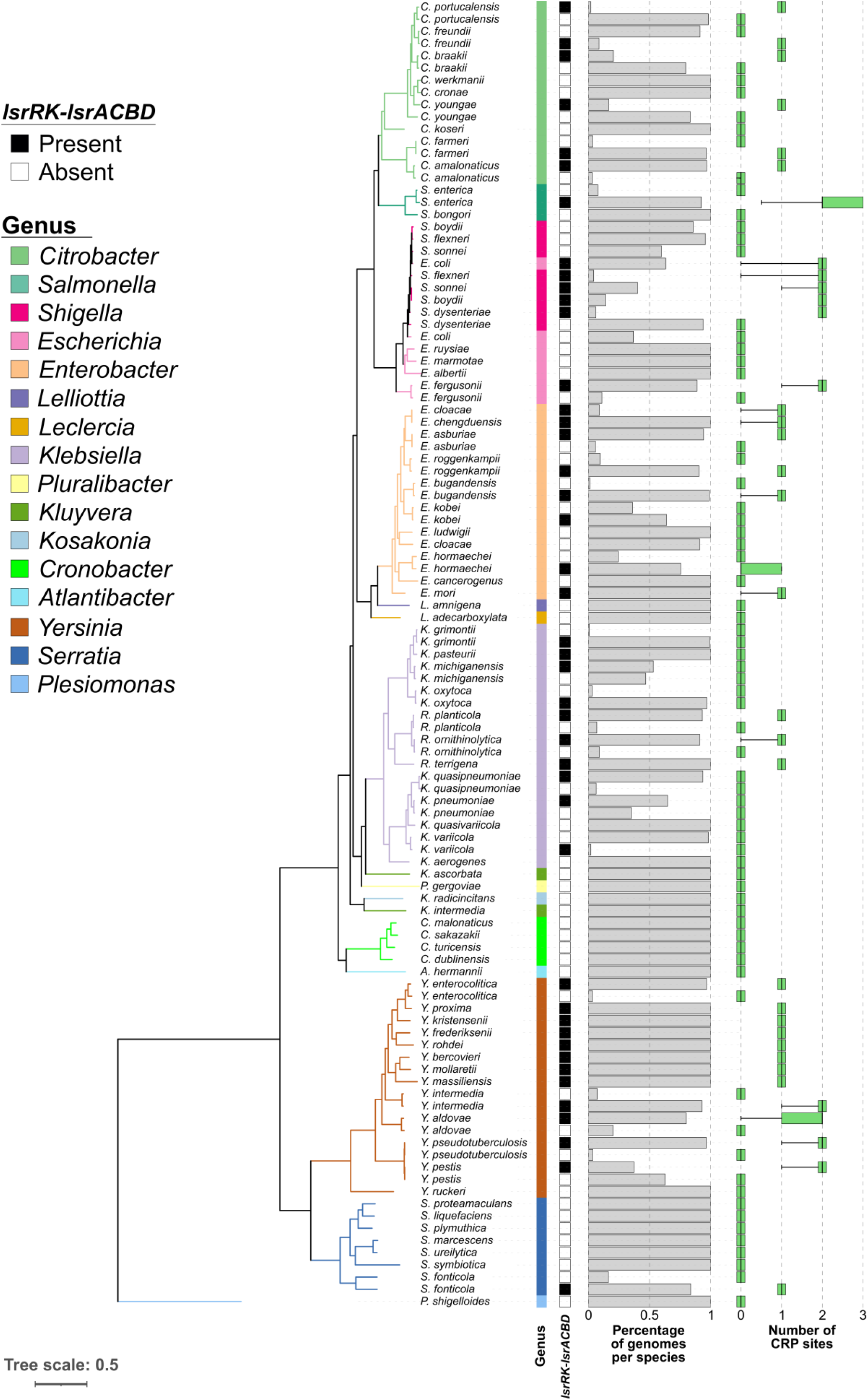
Number of CRP binding sites within the lsr operon across *Enterobacteriaceae*. Phylogenetic relationships between representative *Enterobacteriaceae* species are shown alongside the presence of the *lsr* operon, percentage of genomes containing the operon, and the number of predicted CRP binding sites identified within the operon region. Where available, representative species with and without the *lsr* operon were randomly selected from an enterobacterial core genome tree (53). The number of predicted CRP binding sites is shown as median (black horizontal line), with hinges representing the 25th and 75th percentile. Whiskers extend from the hinges to the maxima and minima, no further than 1.5*distance of the interquartile range. Predicted CRP binding sites vary between 0 to 3 sites across *Enterobacteriaceae*.

Across all genomes containing the *lsr* operon, predicted CRP binding sites were consistently located in proximity to *lsrA* and rarely in intragenic regions (Fig. S3), suggesting a low false positive rate. Approximately 50% of genomes encoding the *lsr* operon (7,610 out of 14,923) harbored at least one CRP binding site in the intergenic region between *lsrA* and i (Fig. 3, Supplementary Table 4). The number of CRP sites per genome ranged from 0 to 3 (mean ± SD = 1.18 ± 1.00), indicating lineage-specific differences in promoter architecture that may reflect variation in the degree of metabolic control over AI-2 responsiveness across Enterobacteriaceae.

### AI-2 import affects intracellular cAMP levels

Having established that *lsr* operon expression is controlled by cAMP-CRP, we next asked whether AI-2 uptake itself influences intracellular cAMP levels, as has been shown for both PTS and non-PTS carbon sources. Intracellular cAMP is sensitive to the PEP/pyruvate ratio: PTS-dependent glucose transport dephosphorylates EIIA^Glc^ and inhibits CyaA, while non-PTS substrates such as maltose and glycerol influence this balance indirectly by altering PEP availability during their metabolism (32).

To monitor intracellular cAMP dynamics in live cells, we employed a FRET-based biosensor consisting of *E. coli* CRP flanked by the fluorophores mTurquoise2 and YFP (36). Upon cAMP binding, CRP undergoes a conformational change that brings the donor and acceptor fluorophores into closer proximity, increasing the YFP/mTurquoise2 emission ratio under donor excitation. A T159A substitution in CRP abolishes DNA binding, ensuring that biosensor expression does not interfere with endogenous CRP-dependent transcription.

To establish a defined low-cAMP baseline, cells were first incubated with glucose prior to sequential addition of AI-2 or non-PTS sugars. As expected, glucose reduced intracellular cAMP levels, reflected by a decreased FRET ratio (Fig. 4A, Fig. S4A). Upon subsequent addition, both AI-2 and all tested non-PTS sugars increased the FRET ratio, indicating elevated cAMP levels. No changes were detected in a Δ*cyaA* mutant (Fig. 4B, Fig. S4B), confirming that the observed signal depends on adenylate cyclase activity rather than a non-specific fluorescence artifact. We note, however, that because glucose establishes a minimal cAMP baseline, any subsequent substrate addition could in principle elevate cAMP levels by relieving glucose-imposed suppression, which limits the specificity of interpretation. Attempts to perform measurements in TB medium without a defined carbon source did not yield interpretable signals, precluding a glucose-independent experimental design (Fig. S5). Finally, deletion of the *lsr* operon did not render the strain insensitive to extracellular AI-2 (Fig. 4C, Fig. S4C). We hypothesize that this is due to the presence of another, yet unidentified, low-affinity AI-2 importer (34). Together, these results indicate that AI-2 uptake, like that of other transported carbon sources, is associated with changes in the intracellular cAMP pool in *E. coli*.

**Fig. 4.**
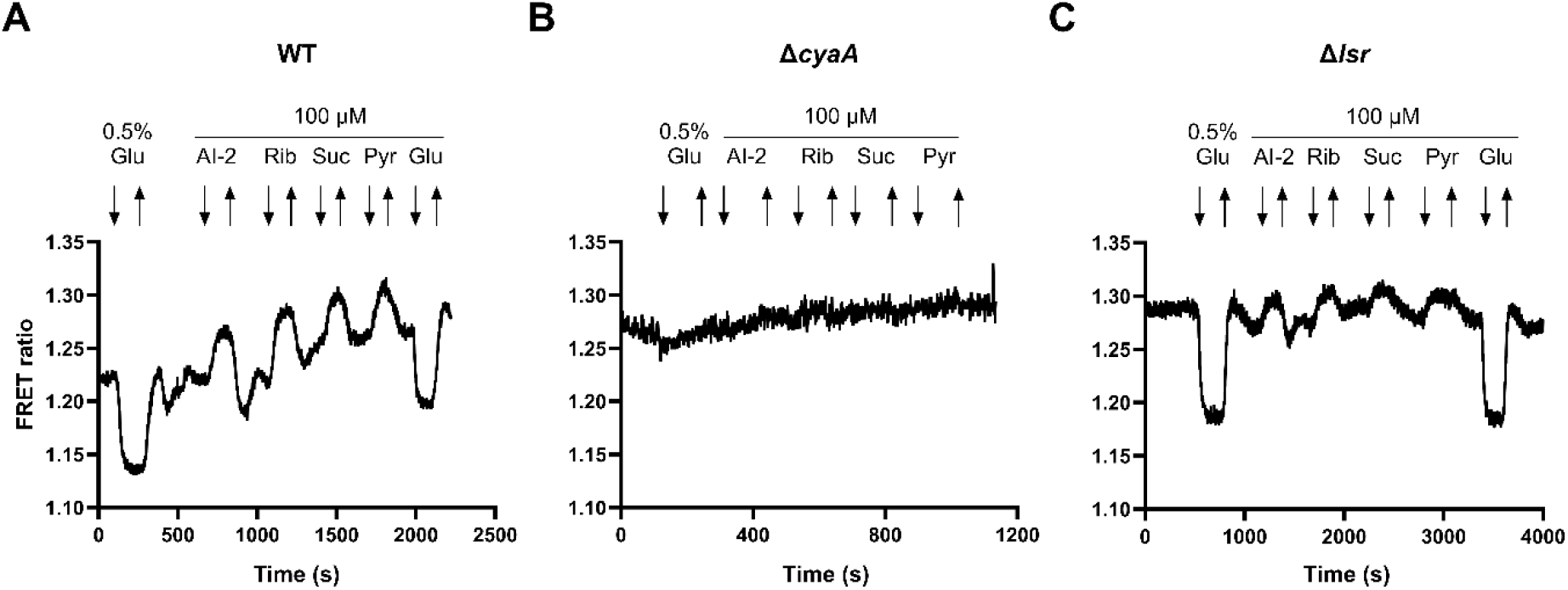
AI-2, non-PTS sugars and gluconeogenic substrates affect cAMP levels in *E. coli*. Representative FRET measurements of intracellular cAMP changes in *E. coli* **(A)** wild-type, **(B)** Δ*cyaA* and **(C)** Δ*lsr* (whole *lsr* operon deletion mutant) strains expressing a plasmid-encoded cAMP sensor. Cells were grown in TB supplemented with 0.2% glucose until mid-exponential phase, harvested and immobilized in a flow chamber, and supplied with constant flow of carbon-free Tanaka medium. After equilibration, cells were stimulated by addition (downward arrow) and subsequent removal (upward arrow) of the indicated carbon sources at specified concentrations. Tested compounds included D-glucose (Glu), autoinducer-2 (AI-2), D-ribose (Rib), L-serine (Ser) and pyruvate (Pyr). FRET signal was obtained by calculating YFP/mTurquoise2 emission ratio under mTurquoise2 excitation.

### AI-2 can serve as carbon source in different bacterial species

Although no examples of *E. coli* strains capable of efficiently utilizing AI-2 as a sole carbon source have been reported (33, 37), the structure and function of the *lsr* operon are remarkably similar to those of canonical sugar utilization operons (Fig. 5A). This makes it a unique case among known quorum-sensing systems. The *lsr* operon encodes an ABC transporter (LsrACDB) that mediates high-affinity uptake of AI-2. Importantly, this system includes the periplasmic binding protein LsrB, which is functionally analogous to binding proteins involved in sugar uptake systems such as maltose (MalE), ribose (RbsB), and galactose (MglB). In these systems, periplasmic binding proteins can also couple ligand binding to chemotaxis signaling via chemoreceptors (38, 39). Similarly to classical sugar operons, the *lsr* operon is transcriptionally repressed by LsrR at low extracellular AI-2 concentrations. Upon uptake, AI-2 is phosphorylated by the kinase LsrK, generating phospho-AI-2, which serves as the intracellular inducer. Phospho-AI-2 binds to LsrR and relieves repression, thereby activating operon expression. This regulatory logic essentially mirrors that of several other sugar utilization systems, although it does not require phosphorylation of the sugar to enable binding to the cognate transcriptional repressor. In addition to uptake and regulation, the *lsr* operon encodes enzymes involved in AI-2 processing and degradation, LsrF and LsrG. These enzymes convert phospho-AI-2 into central metabolic intermediates, including dihydroxyacetone phosphate (DHAP) and acetyl-CoA, linking AI-2 turnover to central metabolism (33). Taken together with our observations that AI-2 import perturbs intracellular cAMP levels and that *lsr* operon expression is controlled by cAMP-CRP, this structural and regulatory parallel raises the possibility that AI-2 once served a primarily metabolic function in *E. coli*, either predating or coexisting with its role as a QS signal. This capacity may have been lost during evolution while signaling function was retained or subsequently acquired.

**Fig. 5.**
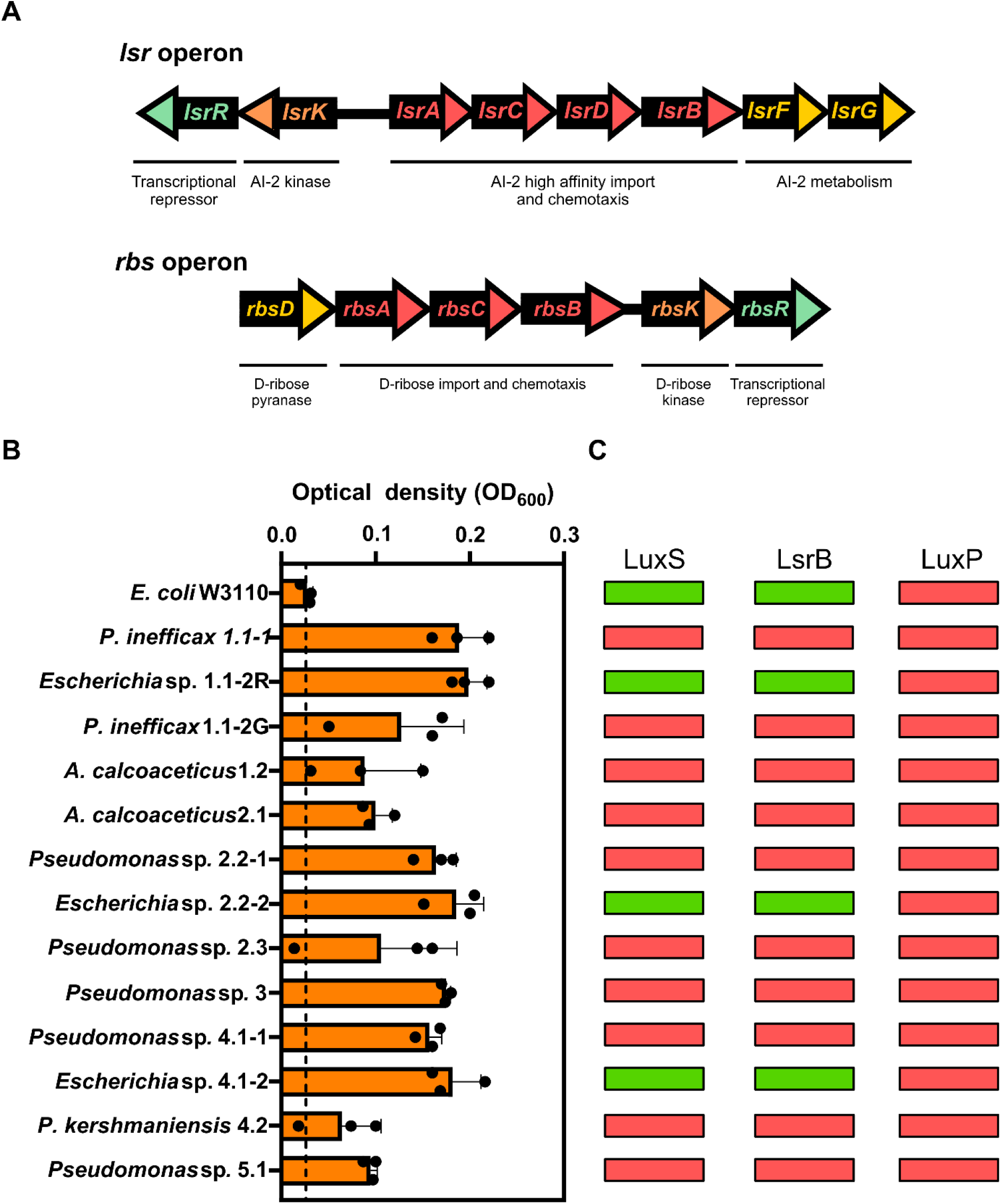
AI-2 can serve as a carbon source in different bacterial species. **(A)** Organization of the AI-2-responsive *lsr* operon and the D-ribose utilization (*rbs*) operon in *E. coli* MG1655. Genes encoding proteins with similar functions are shown in the same color. The sizes of the depicted genes are not proportional to their actual lengths. **(B)** Final optical densities (OD_600_) of cultures of *E. coli* W3110 (negative control) and AI-2-metabolizing isolates grown in M9 minimal medium supplemented with 3 mM DPD/AI-2 as the sole carbon source for 72 h at 30 °C. The dashed line represents the mean OD_600_ value of *E. coli* W3110 culture. Bars indicate mean values (n=3, from two independent experiments), and error bars represent the standard deviation. **(C)** Presence (green) or absence (red) of the AI-2 synthase LuxS, as well as AI-2 receptors LsrB and LuxP, in *E. coli* W3110 and AI-2-metabolizing isolates, based on genome annotation of the isolates with Prokka v1.14.6.

To test whether this metabolic potential persists in environmental bacteria, we collected grass and soil samples from the phyllosphere and rhizosphere, habitats that are associated with metabolically versatile bacteria capable of utilizing a broad range of substrates. Washed suspensions were used to inoculate M9 minimal medium supplemented with 3 mM synthetic DPD/AI-2 as the sole carbon source and incubated statically for 48 h at 30 °C. We observed clear bacterial growth, as indicated by an increase in optical density. The cultures were subsequently streaked onto LB and MacConkey agar plates to isolate and differentiate individual colonies. In total, 14 isolates were obtained, and 13 of these were confirmed to grow reproducibly in M9 minimal medium containing 3 mM DPD/AI-2 as the sole carbon source (Fig. 5B).

Whole-genome sequencing and downstream analysis revealed that the majority of isolates belonged to the genus *Pseudomonas*. In addition, two isolates were identified as *Acinetobacter calcoaceticus*, and two others belonged to the genus *Escherichia* (Fig. 5B). Interestingly, only the *Escherichia* isolates encoded the AI-2-binding protein LsrB, and none of the isolates encoded the alternative AI-2 receptor LuxP (Fig. 5C) (12). Furthermore, neither the *Pseudomonas* nor *Acinetobacter* isolates encoded LuxS, indicating that these organisms do not produce AI-2 themselves and therefore interact with it exclusively as a substrate rather than as a self-produced signal. Whether AI-2 additionally serves a sensory or signaling function in any of these isolates remains unclear. However, the ability to utilize an interspecies signal as a nutrient raises the intriguing possibility that AI-2 catabolism may shape the dynamics of mixed microbial communities, with some members consuming a signal produced by others.

## Discussion

Bacteria inhabit environments that are both spatially and temporally heterogeneous, requiring continuous integration of diverse signals (e.g., nutrients, stress cues, QS autoinducers) to guide their behavior. Rather than operating through isolated pathways, they process this information via interconnected regulatory networks comprising global regulators, two-component systems, small RNAs, and pathway-specific transcription factors, enabling gene expression to reflect the integrated physiological state rather than any single input. (6, 27, 40, 41).

The AI-2 QS system in *E. coli* exemplifies such integration. AI-2 production is coupled to activated methyl cycle activity via LuxS (10), and extracellular AI-2 is imported by the LsrACDB transporter, which additionally mediates chemotaxis through LsrB and Tsr (20). Beyond LsrR-dependent regulation, lsr operon activity is subject to CCR via cAMP-CRP. cAMP-CRP activity depends on the phosphorylation state of EIIA^Glc^, which is coupled to glucose uptake via the PTS and modulated by the intracellular PEP/pyruvate ratio (23, 24, 27, 32). In this study, we built upon existing knowledge of CCR-dependent regulation of the *lsr* operon by systematically testing the effects of both PTS and non-PTS substrates, including sugars, pyruvate, TCA cycle intermediates, and L-serine. Consistent with previous observations, we found that both glucose and non-PTS sugars inhibited *lsr* operon expression, whereas the other tested substrates had no significant effect. This inhibition was dependent on cAMP-CRP and independent of LsrR. We hypothesize that the inhibitory effect of non-PTS sugars may arise from shifts in the PEP/pyruvate ratio, which influence phosphoryl group flux through the PTS and ultimately modulate CyaA activity, thereby reducing *lsr* operon expression. This model may also explain the previously observed inhibition of *lsr* expression in a *ptsI* mutant (42). The *ptsI* gene encodes enzyme I, the first component of the PTS phosphorylation cascade; its absence leads to reduced phosphorylation of EIIA^Glc^, which in turn decreases CyaA activity and limits cAMP-CRP-dependent activation of the *lsr* operon.

Comparative genomic analysis across nearly 20,000 *Enterobacteriaceae* genomes indicates that this regulatory logic is broadly conserved. Around half of the strains encoding the complete *lsr* operon contain CRP binding sites upstream of *lsrA*, with lineage-specific variation in binding site number suggesting differences in regulatory tuning. It is important to note that our motif was derived from experimentally validated sequences in the CollecTF database, which did not include representatives from all genera analyzed in this study. Consequently, divergent CRP motifs in underrepresented or absent genera may not have been captured by our search motif. However, the detection of CRP binding sites in species not represented in the database supports the robustness of our approach and suggests that it can identify such sites beyond the original reference set. CRP binding site architecture near the *lsr* promoter have been maintained across this diverse family, pointing to a possible fitness benefit of coupling QS responsiveness to carbon source availability.

Repression of *lsr* expression by sugars may be further reinforced at the post-translational level. The unphosphorylated form of the PTS phosphocarrier HPr has been shown to directly inhibit LsrK, preventing AI-2 phosphorylation and thereby blocking relief of LsrR-mediated repression (43). This parallels classical inducer exclusion, in which dephosphorylated EIIA^Glc^ inhibits the lactose permease LacY in the presence of glucose, and suggests that metabolic control of the lsr system operates at multiple regulatory levels simultaneously (27).

A particularly intriguing observation from our study is that, similar to other transported substrates, AI-2 uptake is associated with changes in intracellular cAMP levels. Although the experimental constraints of the FRET-based cAMP reporter limit qualitative interpretation of detected changes in cAMP levels, it is notable that AI-2 behaved similarly to non-PTS substrates in these assays. Together with the observation that the *lsr* operon structurally and functionally resembles canonical sugar utilization operons, this raises the possibility that AI-2 is perceived not only as a quorum sensing signal but also as a metabolic input. Interestingly, recently identified mammalian host-derived AI-2 mimics are the keto-pentoses L-xylosone and L-xylulose (44).

This leads to the hypothesis that the *lsr* system may have originally evolved for AI-2 uptake and degradation as a nutrient source, either prior to or alongside its role in quorum sensing. Indeed, intracellular processing of AI-2 yields DHAP and acetyl-CoA, both central metabolic intermediates (33). However, *E. coli* strains tested to date do not grow on AI-2 as a sole carbon source (33, 37), suggesting that under relatively nutrient-rich conditions, AI-2 catabolism may not confer a significant fitness advantage and that this capability may have been lost. In contrast, we were able to isolate soil- and phyllosphere-associated bacteria capable of utilizing AI-2 as a sole carbon source. Interestingly, with the exception of *Escherichia sp*., all isolated bacteria appear to be strict aerobes, suggesting a potential role of oxygen in AI-2 metabolism. The extended lag phases and relatively low final cell densities observed indicate that AI-2 is a comparatively poor substrate. For this reason, we cannot fully assess the extent to which AI-2 contributes to energy generation in *E. coli* when utilized alongside other carbohydrates in nutrient-rich environments such as the mammalian gut. Nevertheless, in nutrient-limited environments such as soil, the ability to metabolize AI-2 may provide a selective advantage. Interestingly, most of these isolates lacked the canonical AI-2 synthase LuxS and did not encode the LsrACDB transporter, suggesting the existence of alternative uptake and degradation pathways. Whether AI-2 plays a signaling role in these organisms remains unclear, but elucidating the underlying metabolic routes represents an important direction for future research.

The apparent paradox that AI-2 is synthesized, secreted, and subsequently reimported for processing or potential metabolism is not unique in bacterial physiology. Similar cycles of overflow and reutilization are well documented for central metabolites. For instance, during rapid growth on excess carbon, *E. coli* excretes acetate as a byproduct of overflow metabolism (45). Upon carbon limitation, however, this acetate is re-assimilated and funneled into central metabolism via acetyl-CoA (46). In this context, AI-2 production and uptake can be viewed as part of a general metabolic logic, where extracellular accumulation followed by reimport allows cells to balance metabolic fluxes while, in case of AI-2, simultaneously enabling additional functions such as intercellular signaling. This could be of physiological importance in growing biofilms, where nutrient-exposed cells produce AI-2 that can be imported and metabolized in nutrient-scarce regions of the biofilm, as previously described for arginine metabolism in *E. coli* biofilms (47).

Taken together, our findings, together with prior work, show that AI-2 regulation integrates specific (AI-2) and global (cAMP) transcriptional regulation. Rather than functioning as an isolated quorum sensing cue, AI-2 appears to be embedded within central metabolic sensing pathways, where its uptake, processing, and regulatory consequences are shaped by the nutritional state of the cells in the population. This positions AI-2 at the intersection of nutrient sensing and population-level communication, suggesting that bacterial decision-making emerges from the convergence of metabolic flux and signaling inputs rather than from dedicated pathways alone. Such integration may also reflect an evolutionary trajectory in which metabolic intermediates were co-opted as communication signals, enabling bacteria to coordinate collective behaviors in a manner that is coupled to the physiological state of the individual cells.

## Materials and Methods

### *E. coli* strains and growth conditions

The strains, plasmids and oligonucleotides used in this study are listed in Supplementary Table 1. All strains were derived from *E. coli* MG1655. Cells were grown either on 1.5% lysogeny broth (LB) agar or in liquid tryptone broth (TB) medium (10 g tryptone and 5 g NaCl per litre), supplemented with kanamycin (50 µg/ml) or ampicillin (100 µg/ml), where necessary. The tested substrates were prepared as sterile 100 mM stock solutions in distilled water. Gene deletions were constructed by PCR-based inactivation of chromosomal genes using λ-Red recombinase and oligonucleotides listed in Supplementary Table 2 (48, 49). Kanamycin resistance cassettes were subsequently removed via FLP recombination (50).

### Flow cytometry

To assess *lsr* operon activity, *E. coli* MG1655 cells were transformed with a plasmid-based GFP reporter containing the 217-nucleotide region upstream of the *lsrA* gene (pVS1723) (17). Overnight cultures grown in TB medium were diluted 1:100 into 3 ml of fresh TB medium and incubated for 3 h at 37 °C with shaking (200 rpm). The tested substrates and synthetic DPD/AI-2 (provided by Dr. Rita Ventura, ITQB, Oeiras, Portugal) (51) were then added to a final concentration of 1 mM and 50 or 100 µM, respectively, followed by an additional 1 h incubation at 37 °C with shaking. Subsequently, cultures were diluted 1:400 into 2 ml PBS (137 mM NaCl, 2.7 mM KCl, 10 mM Na_2_HPO_4_, 1.8 mM KH_2_PO_4_), and fluorescence was measured using a BD LSRFortessa SORP cell analyzer (BD Biosciences, Germany).

### Complementation of *crp* deletion

To complement the Δ*crp* mutation, the wild-type *crp* allele was amplified from the *E. coli* MG1655 chromosome using primers listed in Supplementary Table 2. The PCR product was cloned into NcoI- and HindIII-digested pTrc99a expression vector downstream of the isopropyl β-D-1-thiogalactopyranoside (IPTG)-inducible promoter using Gibson assembly (Gibson Assembly^®^ Master Mix, NEB, USA) according to the manufacturer’s instructions. The resulting pTrc99a::*crp* construct was introduced into *E. coli* MG1655 Δ*crp* strain, and IPTG titration was performed to determine the expression level that restored wild-type-like regulation of the *lsr* operon, as measured by flow cytometry. The optimal IPTG concentration (7.5 µM) identified in this assay was used in all subsequent experiments.

### Plasmid-based cAMP sensor construction and FRET measurements

To construct cAMP FRET sensor, pTrc99a plasmid was linearized by NcoI and HindIII restriction enzyme digestion. Genes encoding YFP, CRP^T159A^, and mTurquoise2 were amplified and assembled into the vector backbone using Gibson assembly with primers listed in Supplementary Table 2, resulting in pXC2 plasmid. Flexible linkers with 5 consecutive glycine residues (GGGGG) were introduced between fusion protein domains.

*E. coli* strains harboring pXC2 plasmid were grown overnight in TB supplemented with 0.2% glucose at 37 °C with shaking (200 rpm). Overnight cultures were diluted 1:200 in fresh medium supplemented with 100 µM IPTG to induce sensor expression. Cells were cultivated to an OD_600_ of ∼0.8, harvested by centrifugation at 4000 rpm for 5 min, and washed twice with Tanaka medium (20 mM (NH_4_)_2_SO_4_, 34 mM Na_2_HPO_4_, 0.3 mM MgSO_4_, 64 mM KH_2_PO_4_, 10 μM CaCl_2_, 1 μM FeSO_4_, and 1 μM ZnCl_2_). Cell suspension was then stored at 4 °C for at least 1 hour prior to measurements.

For microscopy, cells were immobilized on a poly-L-lysine-coated coverslip for 10 min, which was subsequently mounted in a flow-through chamber and subjected to a constant flow (0.3 mL/min) of Tanaka medium using a syringe pump (Harvard Apparatus, USA). Cells were first equilibrated in Tanaka media and then exposed to the addition or removal of specific carbon sources. Measurements were performed as previously described using a Zeiss Axio Imager.Z1 microscope equipped with a 40x/0.75 objective and a photon counting detector (Hamamatsu, Japan) (52). A dense monolayer of cells expressing the cAMP sensor were excited at 436/20 nm. Emission from YFP (520 nm long-pass) and mTurquoise2 (480/40 nm band-pass) emissions was recorded simultaneously with an integration time of 1 s.

Time-series fluorescence intensities were corrected for photobleaching using MATLAB R2020a (MathWorks, USA) as described before (36). For individual trace, fluorescence decay was fitted with an exponential decay (*f*(*t*) = *ae*^−*bt*^ + *c*), where *f(t)* is the fluorescence intensity at time *t, a* is the amplitude, *b* is the bleaching rate constant, and *c* is the offset. Fitting was performed separately for each channel using only the baseline preceding stimulation and then the fitted decay was extrapolated over the entire duration. The raw YFP and mTurquoise2 fluorescence traces were then corrected using corresponding fitted bleaching curves, and the corrected signals were subsequently used to calculate FRET ratio.

### Analysis of CRP binding sites in *Enterobacteriaceae* genomes

#### Genome dataset and identification of the lsr operon

A collection of 19,795 Enterobacteriaceae genomes was obtained from a study by Näpflin and Schubert et al (53). Briefly, high-quality genomes classified as *Enterobacteriaceae* or *Yersinia* were obtained from ProGenomes v3 (54). Species represented by less than 10 genomes were discarded, and overrepresented species were randomly subset resulting in a genome collection spanning 16 genera and 80 species (Supplementary Table 3).

To identify the *lsr* operon across *Enterobacteriaceae* and *Yersinia* genomes, predicted protein sequences were screened using hmmscan (HMMER v3.1b2) with hidden Markov models (HMM) profiles constructed for each *lsr* operon protein (55). Protein-coding sequences were predicted using Prodigal v2.6.3 (56). For HMM construction, reference sequences corresponding to each operon gene (*lsrK, lsrR, lsrA, lsrC, lsrD, lsrB, lsrF, lsrG*, and *tam*) were downloaded from UniProt (accessed 02/06/2026), and sequences not originating from *Enterobacteriaceae* (NCBI taxon id: 543) were excluded (57). Filtered sequences were aligned with mafft v7.515 (--maxiterate 1000 --localpair) and HMM profiles were built using hmmbuild (HMMER v3.1b2) (55, 58). Potential homologs were retained using E-value < 1e-20 and a protein length within ± 50% of the corresponding *E. coli* reference sequence. Putative *lsr* operons were only retained if *lsrA* and *lsrR* were found in directly adjacent genomic locations. Genomic coordinates, strand orientation, and operon completeness were recorded for downstream analyses. For each genome containing the *lsr* locus (containing at least the complete region from *lsrK* to *lsrB*), the genomic region encompassing the complete *lsr* operon together with 1000 bp upstream and downstream flanking regions was extracted and used as input for motif scanning.

#### Motif definition and scanning for CRP binding sites

CRP binding motifs were represented as position weight matrices (PWMs) derived from the collecTF, a curated database for promotors of different species (59). We downloaded all available motifs from http://collectf.org/browse/search_terms/?search-term=crp (accessed online 03/2026) and converted them into MEME format to use it for genome-wide motif scanning.

The motif scanning was performed using FIMO, applying a stringent significance threshold (*p* < 6.1e−5) (60). A precomputed background model (Markov model) derived from all our genomic sequences was used to improve the specificity. FIMO reports overlapping and highly redundant motif hits, thus hits were deduplicated to only report the best hit of multiple overlaps (Supplementary Table 4).

Next, all motif hits were filtered for distance from *lsrA* and *lsrR*, to ensure that the CRP sites are selected from the correct region in between the two genes. When not applying this filtering to *lsrR* (e.g. also allow “erroneous” hits on the opposite of *lsrA* or in intragenic regions) a false positive rate of approximately 0.0035% could be estimated. To visualize the distribution of the presence of the *lsr* operon and the number of CRP binding sites across the phylogeny of *Enterobacteriaceae*, the data was mapped onto the core genome phylogenetic tree provided by Näpflin and Schubert et al (53). The phylogenetic tree was randomly subset to include a random species with and without the *lsr* operon present, where available, as a representative using ete3 toolkit v 3.1.3 (61). ITOL v7.5.1 was used for tree visualization and the number of CRP binding sites were visualized using boxplots for each species (62).

### Isolation of AI-2-metabolizing bacterial isolates

To isolate bacteria capable of utilizing AI-2 as a sole carbon source, 3 grass and 2 soil samples were collected on the Hönggerberg campus of ETH Zurich (Zurich, Switzerland). The samples were washed in sterile PBS and inoculated at a 1:100 dilution into 1 ml of M9 minimal medium (6.8 g Na_2_HPO_4_, 3 g KH_2_PO_4_, 0.5 g NaCl, 1 g NH_4_Cl, 0.24 g MgSO_4_, 0.011 g CaCl_2_ per litre) supplemented with 3 mM DPD/AI-2, followed by incubation without shaking for 48 h at 30 °C. Bacterial growth was detected in all samples, and 5 µl of each culture was streaked onto LB and MacConkey agar plates to enable phenotypic differentiation of the isolates. A total of 14 isolates were obtained and subsequently tested for their ability to grow in M9 minimal medium using AI-2 as the sole carbon source.

Overnight cultures of the isolates were grown at 30 °C with shaking (200 rpm) in M9 minimal medium supplemented with 1% glucose or arabinose (for isolates 1.2 and 2.1, later identified as *A. calcoaceticus* strains). The cultures were then washed in PBS and inoculated into M9 minimal medium supplemented with 3 mM DPD/AI-2, followed by incubation for 72 h without shaking at 30 °C. Growth was assessed by measuring OD_600_ using an Infinite 200Pro plate reader (Tecan, Switzerland). Growth in M9 minimal medium containing 3 mM DPD/AI-2 was confirmed for 13 of the 14 isolates, which were subsequently used for further analysis.

### Identification and bioinformatic analysis of AI-2-metabolizing bacterial isolates

Genomic DNA was extracted from pure bacterial cultures using QIAamp DNA Extraction Kit (QIAGEN, Germany) according to the manufacturer’s instructions. Sequencing libraries were prepared by the NGS core facility of the MPI for Biochemistry (RRID: SCR_025746) using NEBnext Ultra II FS DNA library prep (NEB, USA) and sequenced on an the Aviti system sequencing instrument (Element Biosciences, USA) to generate paired-end FASTQ files with 2 x 75 bp reads. Read quality was assessed with FastQC (https://www.bioinformatics.babraham.ac.uk/projects/fastqc/). Adapter content and per-base quality profiles were inspected before assembly.

Paired-end sequencing reads were *de novo* assembled with Unicycler (v. 0.5.1) using the internal SPAdes algorithm (v. 4.1.0) as the *de novo* assembler (63, 64). Assemblies were then run in normal mode (`--mode normal --keep 3 --verbosity 2`). For exploratory assemblies of concatenated first-run and resequencing reads, Unicycler was run with a fixed k-mer set (`--kmers 13,25,33,39,45,49,53,57`) to account for the different read lengths between sequencing runs. The retained assembly FASTA files were `assembly.fasta` files generated for each sample. Genome annotation was performed with Prokka v1.14.6. For each isolate, Prokka was run with an e-value threshold of `1e-9` (65).

Taxonomic classification was performed from the assembled contigs using Kraken2 with the standard Kraken2 database `k2_standard` (v. 20250402) (66). For each isolate, Kraken2 was run against the retained `assembly.fasta` file, producing both per-contig classification output and a taxonomic report. Kraken2 reports were further processed with Bracken using the same database, a read length parameter of 150 (`-r 150`), and a minimum count threshold of 10 (`-t 10`) to estimate taxon-level abundance from the Kraken2 classifications (67). Kraken2 and Bracken reports were used to assign the taxonomic identity of each isolate. In cases of inconsistency between Kraken2 and Bracken outputs, classification of the isolate was limited to the genus level.

Genes of interest were screened in the Prokka annotation outputs. Annotation files were searched for `luxS`, `lsrB`, and `luxP`, corresponding to S-ribosylhomocysteine lyase LuxS, AI-2-binding periplasmic protein LsrB, and AI-2-binding periplasmic protein LuxP, respectively.

## Acknowledgements

The authors thank Moritz Klaßmann for the construction of *E. coli* MG1655 Δ*lsr* strain and Dr. Yu Chen for the construction of *E. coli* MG1655 Δ*cyaA* strain. This work was supported by grant LA 4572/4-1 from the Deutsche Forschungsgemeinschaft.

## Contributions

L.L. and C.S. conceived and designed the study. N.G. and X.C. performed the experiments. N.N., L.M., and A.Y. performed the bioinformatics analysis. L.L., N.G., and C.S. wrote the manuscript with contributions from all authors.

## Code and data availability

All code and a Nextflow pipeline to identify the *lsr* operon and predict CRP binding sites can be obtained under https://github.com/NicolasNaepflin/lsr_crp_finder and https://doi.org/10.5281/zenodo.20407787. The whole-genome sequencing data of the AI-2-metabolizing isolates are available on NCBI under accession numbers SAMN57324306– SAMN57324318.

