## Supplementary Figures 1-5 for "Autoinducer-2 functions as both a quorum sensing and metabolic signal in *Escherichia coli*"

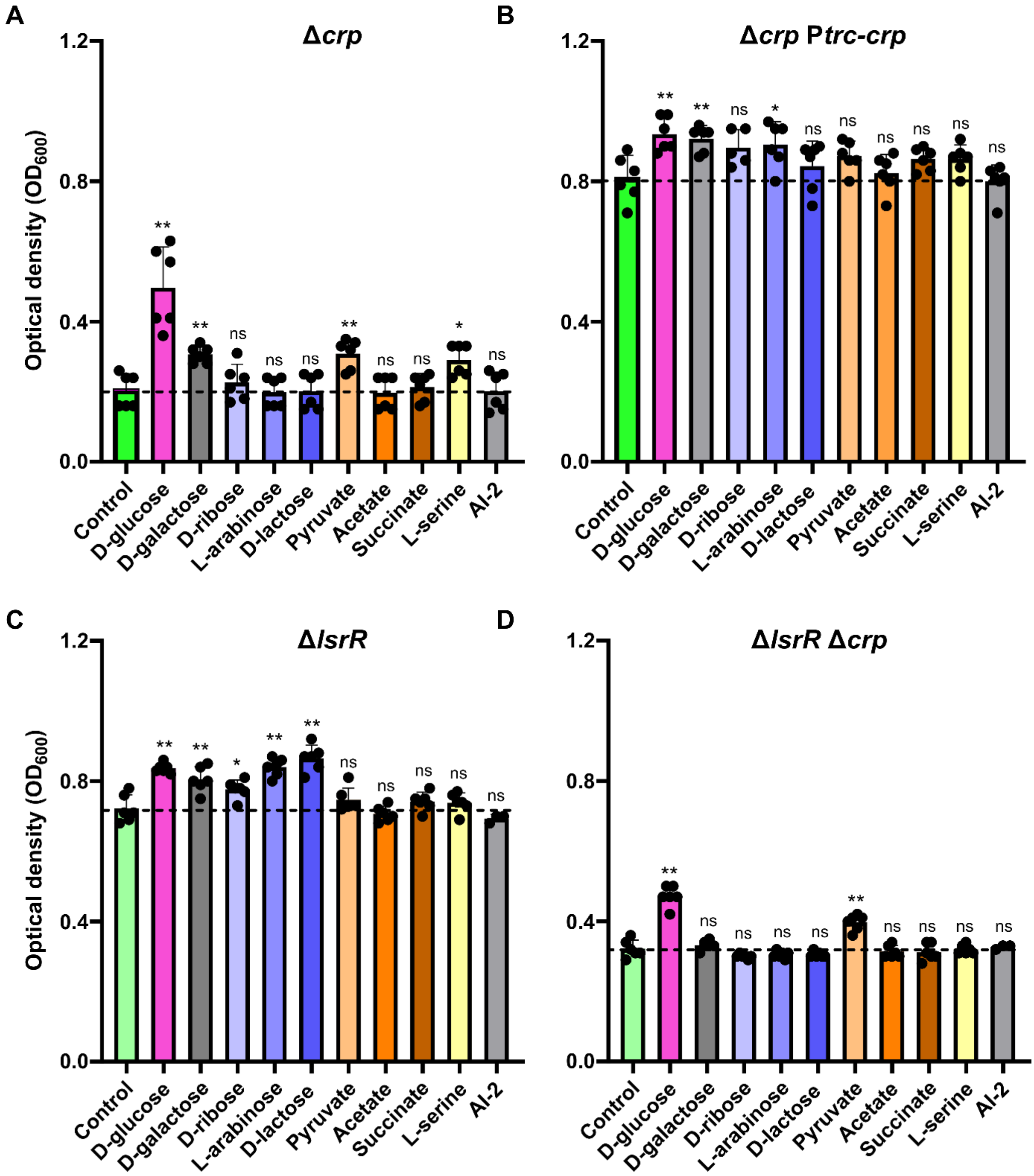


**Fig. S1. Optical densities of the cultures shown in Fig. 2A-D.** The dashed line represents the mean value of the control. *P* values were calculated using a two-tailed Mann-Whitney *U*-test (**P<0.01; *P<0.05; ns, not significant). Bars indicate mean values (n=6, from two independent experiments), and error bars represent the standard deviation.

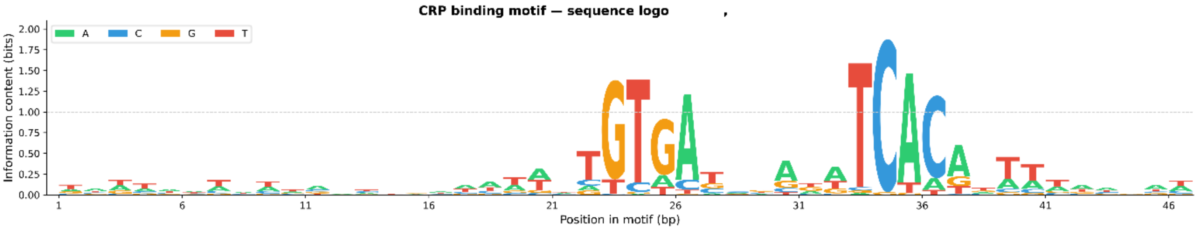


**Fig. S2. CRP binding motif in *Enterobacteriaceae.*** Sequence logo of the CRP binding motif generated from 119 unique CRP binding sites detected within the *lsr* operon in *Enterobacteriaceae,* obtained from CollecTF*.* Conserved nucleotides are found at positions 23-26 and 33-36.

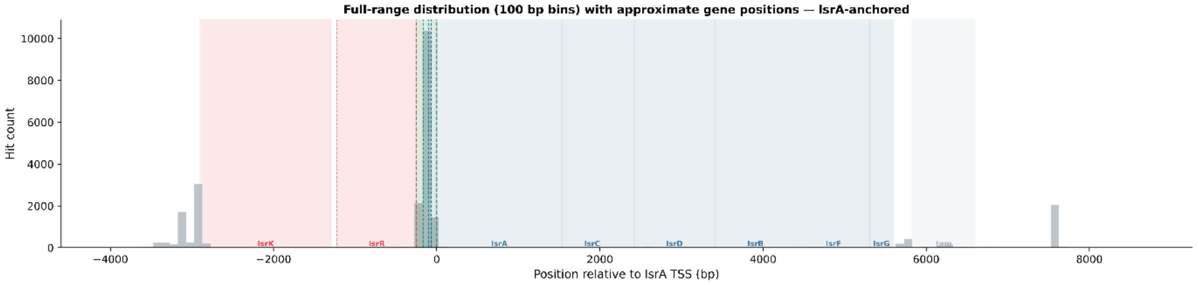


**Fig. S3. Distribution of CRP binding sites within the *lsr* operon and its genomic context.** Binding sites are shown relative to the *lsrA* transcription start site (TSS), with hits grouped into 100 bp bins. Boxes indicate approximate gene boundaries of genes within the *lsr* operon. The strongest enrichment of CRP binding sites is located between the genes *lsrA* and *lsrR*.

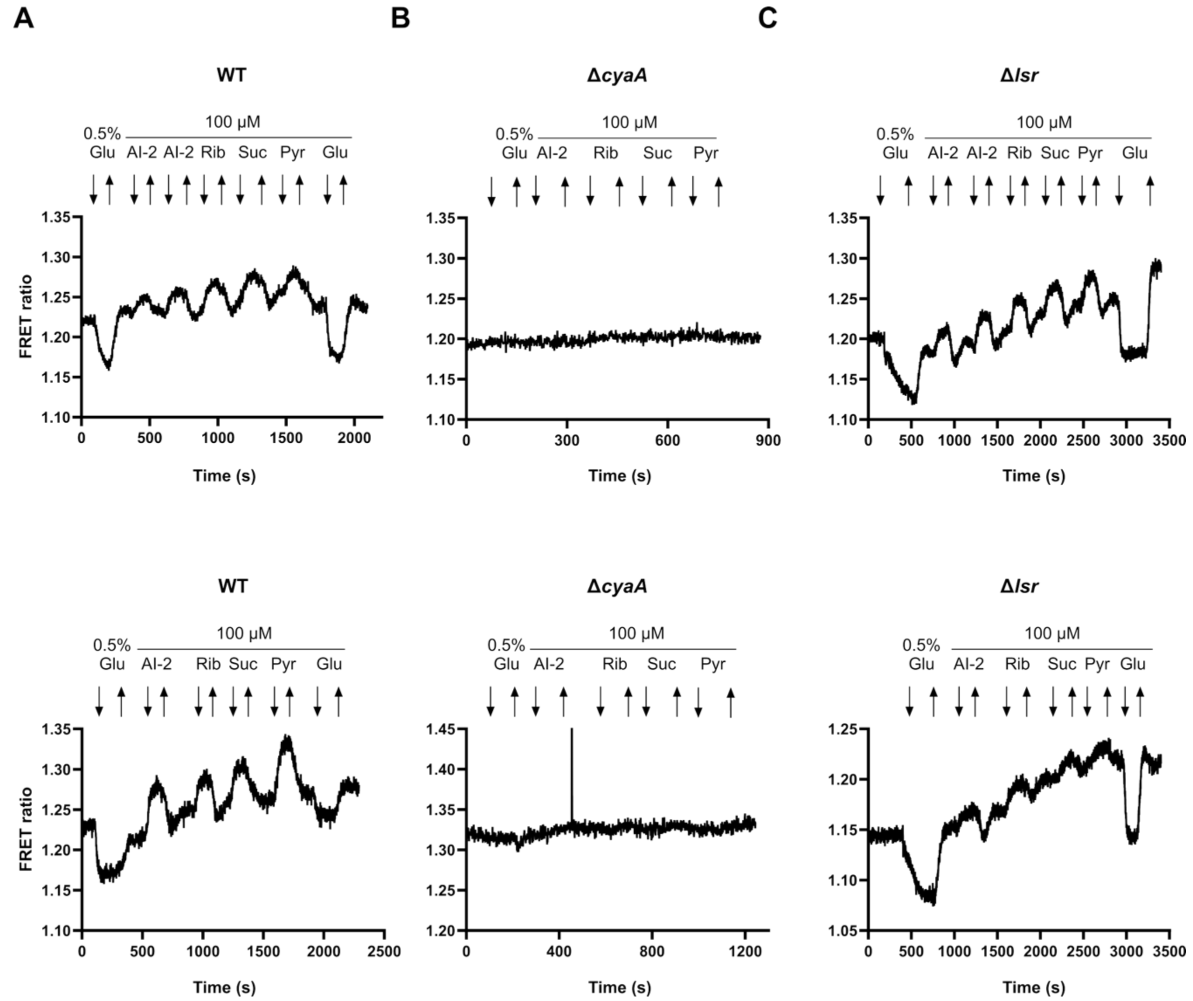


**Fig. S4. Replicates of FRET measurements confirm cAMP response in *E. coli*, corresponding to Figure 3.** Independent biological replicates of the experiments shown in Figure 3 performed under the same conditions.

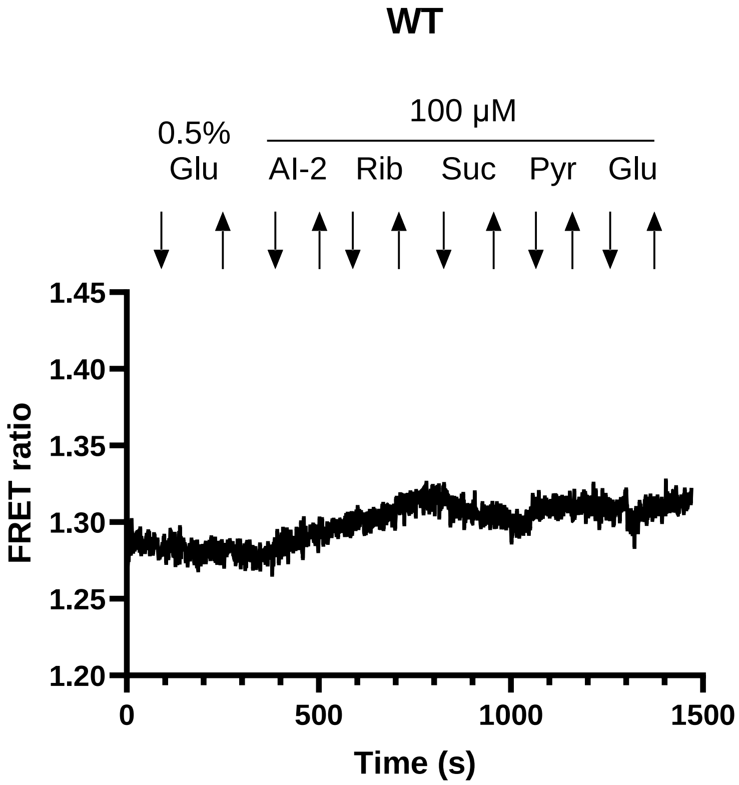


**Fig. S5.** Representative FRET measurements of intracellular cAMP changes in *E. coli* grown in TB medium in absence of glucose. Cells were grown in TB until mid-exponential phase, harvested and immobilized in a flow chamber, and supplied with constant flow of carbon-free Tanaka medium. After equilibration, cells were stimulated by addition (downward arrow) and subsequent removal (upward arrow) of the indicated carbon sources at specified concentrations. Tested compounds included D-glucose (Glu), autoinducer-2 (AI-2), D-ribose (Rib), succinate (Suc) and pyruvate (Pyr). FRET signal was obtained by calculating YFP/mTurquoise2 emission ratio under mTurquoise2 excitation.
